# Reticulon and CLIMP-63 regulate nanodomain organization of peripheral ER tubules

**DOI:** 10.1101/550715

**Authors:** Guang Gao, Chengjia Zhu, Emma Liu, Ivan R. Nabi

## Abstract

The endoplasmic reticulum (ER) is an expansive, membrane-enclosed organelle composed of smooth peripheral tubules and rough, ribosome-studded central ER sheets whose morphology is determined, in part, by the ER-shaping proteins, reticulon and CLIMP-63, respectively. Here, STimulated Emission Depletion (STED) super-resolution microscopy shows that reticulon4a (RTN4a) and CLIMP-63 also regulate the organization and dynamics of peripheral ER tubule nanodomains. STED imaging shows that lumenal ERmoxGFP, membrane Sec61βGFP, knock-in calreticulin-GFP and antibody-labeled ER resident proteins calnexin and derlin-1 are all localized to periodic puncta along the length of peripheral ER tubules that are not readily observable by diffraction limited confocal microscopy. RTN4a segregates away from and restricts lumenal blob length while CLIMP-63 associates with and increases lumenal blob length. RTN4a and CLIMP-63 also regulate the nanodomain distribution of ER resident proteins, being required for the preferential segregation of calnexin and derlin-1 puncta away from lumenal ERmoxGFP blobs. High-speed (40 ms/frame) live cell STED imaging shows that RTN4a and CLIMP-63 regulate dynamic nanoscale lumenal compartmentalization along peripheral ER tubules. RTN4a enhances and CLIMP-63 disrupts the local accumulation of lumenal ERmoxGFP at spatially defined sites along ER tubules. The ER shaping proteins reticulon and CLIMP-63 therefore regulate lumenal ER nanodomain heterogeneity, interaction with ER resident proteins and dynamics in peripheral ER tubules.

## Introduction

Since the initial Singer-Nicholson fluid mosaic model of free membrane diffusion in the 80’s, the role of membrane nanodomains in the control of protein and lipid dynamics in the plasma membrane, regulating signal transduction, endocytosis and exocytosis and thereby cellular behavior, has been extensively characterized (1). In contrast, the nanodomain organization of organellar membrane structures remains poorly defined.

The endoplasmic reticulum (ER) is a continuous membrane network that is classically divided into central ribosome-studded rough ER sheets, the site of protein synthesis, and peripheral smooth ER tubules, implicated in lipid synthesis and detoxification (2–5). A family of ER shaping proteins that maintain sheet or tubule architecture include the cytoskeleton-linking membrane protein 63 (CLIMP-63), ribosome-interacting protein p180, reticulon (RTN), atlastin (ATL), and DP1/Yop1p (4–8). RTN has two hydrophobic hairpins which could form a wedge-like structure, causing local ER curvature by replacing the lipids in the outer leaflet of the lipid bilayer (9) and has been implicated in peripheral ER tubule formation (3, 10–12). CLIMP-63 is predominantly associated with central ER sheet formation and has been proposed to function as a spacer that maintains ER sheets (6). Atlastin induces membrane fusion and the formation of 3-way junctions (4, 8). Reticulon family member RTN4a induces expansion of peripheral ER tubules while CLIMP-63 promotes ER sheet formation; the relative expression of these two ER shaping proteins determines the cellular abundance of ER sheets versus tubules (3, 6, 12). CLIMP-63 has also been localized to peripheral ER tubules (13) and, here, we use STED super-resolution imaging to show that RTN4a and CLIMP-63 regulate the nanodomain organization and dynamics of peripheral ER tubules.

The thickness of an ER sheet and the diameter of an ER tubule is typically 30-100 nm (2, 14), below the diffraction limit of visible light (~200 nm), hindering the characterization of ER structure by standard confocal fluorescence microscopy. In addition to these spatial limitations, the ER is a highly dynamic organelle and its study in live cells requires high temporal resolution; recent use of high-speed, super-resolution imaging techniques suggested that peripheral ER sheets are densely packed tubular arrays (13). STED super-resolution microscopy (15) obtains lateral resolution of ~50 nm and the pioneering application of STED to the ER revealed ring structures formed by the tubular network of the ER that were not observed by conventional confocal microscopy (16). A more recent STED analysis identified the presence of dynamic nanoholes in peripheral ER sheets (17). Single molecule super-resolution particle tracking of an ER lumenal reporter demonstrated the existence of active ER lumenal flow in peripheral ER tubules (18) while high-speed, super resolution GI-SIM microscopy identified lumenal bulges and constrictions along ER tubules (19).

However, molecular mechanisms that regulate organization of ER nanodomains remain to be defined. Here we apply STED microscopy to study lumenal compartmentalization in peripheral ER tubules using the lumenal ER reporter ERmoxGFP. ERmoxGFP contains the bovine prolactin signal sequence and KDEL ER retention sequence linked to monomeric, cysteine-less moxGFP, a modified inert GFP optimized for use in oxidizing environments that minimally perturbs the cell (20). We find that ERmoxGFP defines local lumenal filling of nanodomains that are segregated from membrane-associated ER proteins along peripheral ER tubules. Reticulon enhances and CLIMP-63 disrupts the spatial localization of these lumenal nanodomains along ER tubules thereby impacting the segregation of membrane-associated ER proteins from lumenal ER nanodomains.

## Results

### STED super-resolution microscopy reveals nanoscale periodicity in ER tubules

The peripheral ER imaged by diffraction limited confocal microscopy presents a highly reticular network of interconnected ER tubules. Super-resolution 2D STED live cell imaging of HT-1080 fibrosarcoma and COS-7 cells transfected with the ER lumenal reporter ERmoxGFP (20) shows that peripheral ER tubules are highly periodic and composed of tubules showing discrete densities of ERmoxGFP (Fig. 1A). Peripheral ER tubule periodicities are observed by 2D STED live cell imaging of ERmoxGFP transfected HT-1080 and COS-7 cells, at a temporal resolution of 0.8 seconds per frame (Supp. Video 1). Peripheral ER tubule periodicities are also observed in cells fixed with 3% paraformaldehyde/0.2% glutaraldehyde (Fig. 1A), an established fixation protocol that preserves ER architecture (13, 21–23).

**Fig. 1.**
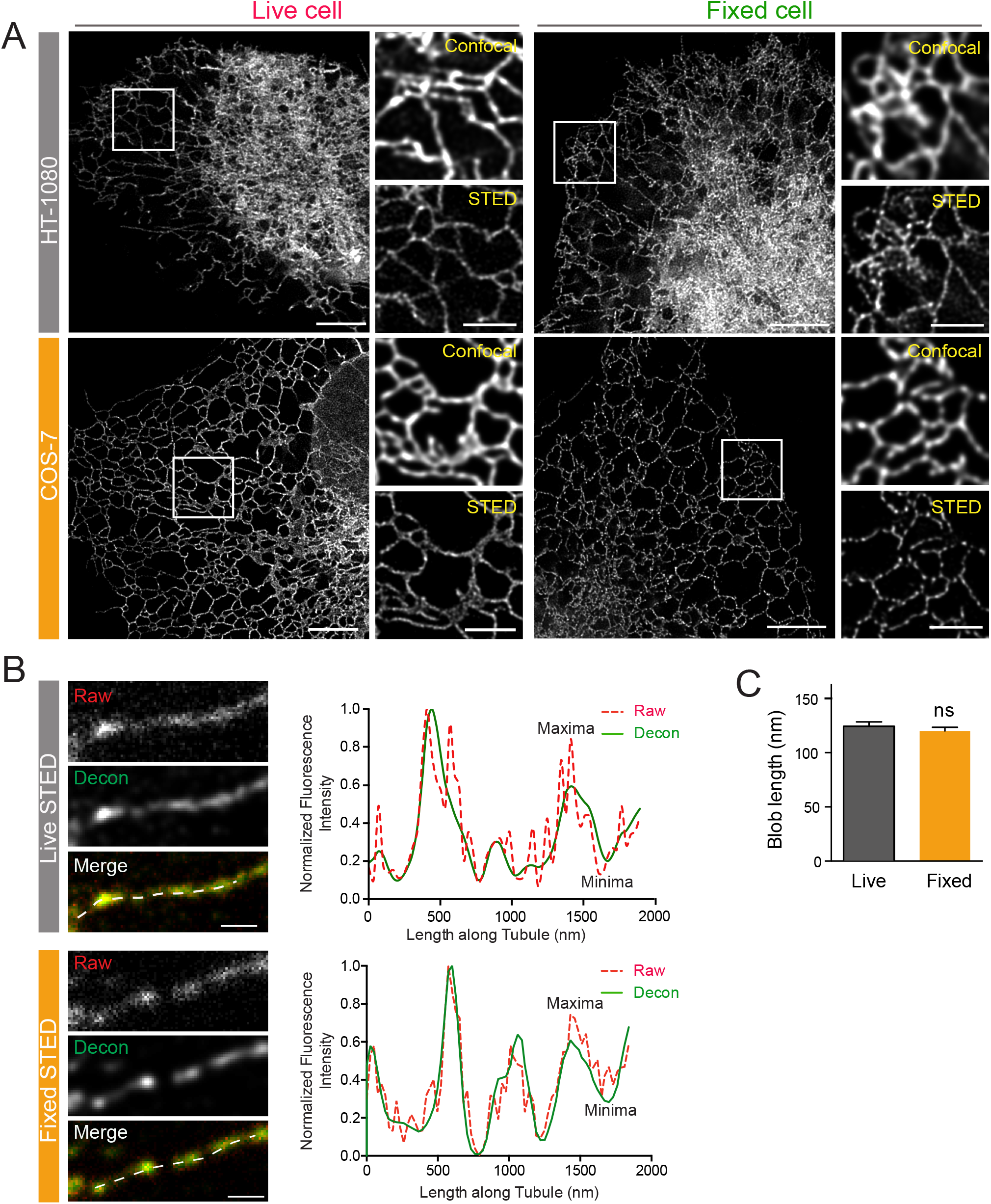
STED imaging reveals lumenal nanodomain periodicity in peripheral ER tubules. A) Representative 2D STED images of live and fixed HT-1080 or COS-7 cells expressing ERmoxGFP are shown. Magnified confocal and STED images of the boxed region highlight the improved resolution obtained by 2D STED. Scale bar: 5 μm; zooms: 2 μm. B) Line scans of isolated peripheral ERmoxGFP labeled tubules (dashed) from STED images of live and fixed cells show matching maxima and minima for raw (red, dashed) and deconvolved (decon, green, solid) images. Scale bar: 0.5 um. C) Length (FWHM) of ERmoxGFP blobs was measured in 2D STED images of peripheral ER tubules in live and fixed cells. Values plotted are mean ± SEM from three independent experiments (20 line scans/each repeat). Significance assessed by student’s t-test. ns, not significant.

Line scan analysis shows distinct maxima and minima, corresponding to local enrichment and depletion, respectively, of fluorescent signal from the lumenal ERmoxGFP reporter along peripheral ER tubules in fixed and live cells (Fig. 1B). ER tubule line scan analysis of raw STED images matches that of the deconvolved image; deconvolution effectively reduces noise along the line scan, that is increased for live cell scans due to refractive index mismatching of the imaging media, without impacting blob size or distribution (Fig. 1B). ER lumenal blob length, determined from the full width at half maximum (FWHM) measurement of maxima in line scan analysis, is equivalent along ER tubules in both live and fixed cells (Fig. 1C). STED analysis of fixed cells has, therefore, retained nanodomain features of ER tubules observed in live cells.

As RTN4a promotes ER tubule formation (3, 10–12), we therefore assessed whether it also regulated peripheral ER tubule periodicity. Upon RTN4 siRNA knockdown, lumenal ERmoxGFP labeled maxima in peripheral ER tubules are elongated relative to control (Fig. 2A,B); CLIMP-63 siRNA knockdown did not alter the periodic distribution of this lumenal ER reporter (Fig. 2A,B). To quantify peripheral ER tubule periodicity, we measured maxima (blob) length, variation in maxima length (SD) and maxima-to-minima fluorescence intensity differentials from line scans of ER tubules (i.e. Fig. 1B). Significantly increased maxima length, increased variation of maxima length and reduction in maxima to minima intensity differential are observed for ER tubules of RTN4 knockdown cells (Fig. 2C). No significant changes in these parameters were observed for ER tubules of CLIMP-63 knockdown cells (Fig. 2C) compared to cells transfected with control siRNA. This suggests that RTN4 not only induces peripheral ER tubule formation but also regulates the nanodomain organization of these ER tubules.

**Fig. 2.**
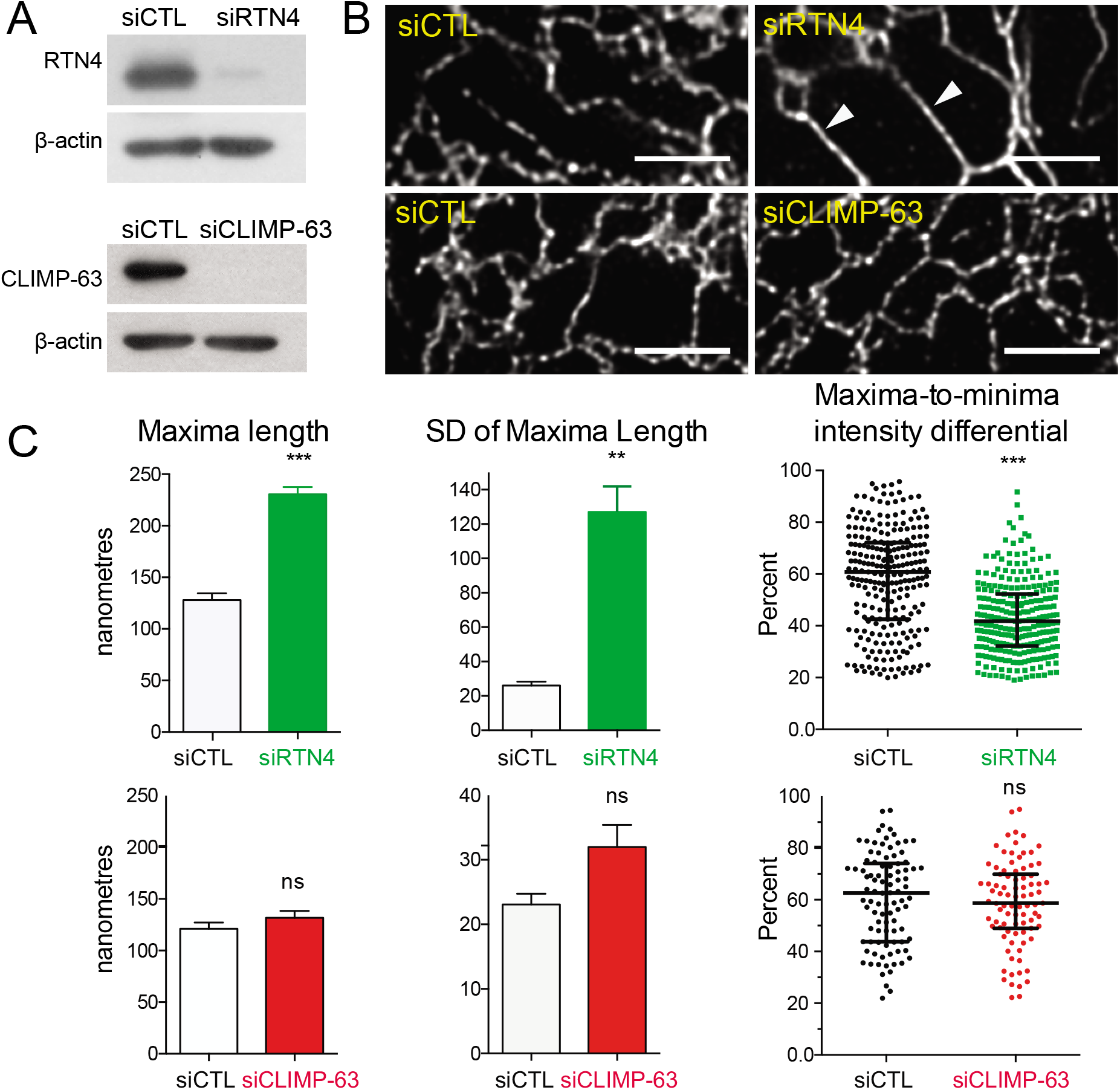
Reticulon regulates lumenal ER nanodomain periodicity. A) Western blots of siCTL, siRTN4, and siCLIMP-63 transfected HT-1080 cells were probed with anti-RTN4a, anti-CLIMP-63, and anti-β-actin as a loading control. B) Representative images of ER tubules in HT-1080 cells transfected with ERmoxGFP and siCTL, siRTN4 or siCLIMP-63. Arrowheads indicate the tubules with increased blob length. Scale bar: 2 μm. C) The quantification of maxima length, variation of maxima length (SD) and maxima-minima intensity differentials of ER tubules in HT-1080 cells transfected with siRTN4, siCLIMP-63 or siCTL. Bar graphs show mean±SEM and scatter dot plots median with interquartile range. Significance assessed by student’s t-test from three independent experiments (40 line scans/each repeat). **, P < 0.01; ***, P < 0.001, ns, not significant.

We then tested whether overexpression of mCherry-RTN4a, mCherry-CLIMP-63 and, as a control, mCherry-ATL1, impacts lumenal nanodomain organization in peripheral ER tubules. ERmoxGFP-transfected HT-1080 cells show expansion of the central ER and reduction of peripheral tubules upon mCherry-CLIMP-63 overexpression, formation of an extended network of peripheral tubules upon mCherry-RTN4a overexpression and increased branched peripheral ER structures upon mCherry-ATL1 overexpression, consistent with previous reports on these ER shaping proteins (3, 6, 9, 12, 24). Similar to RTN4 knockdown (Fig. 2 B,C), mCherry-CLIMP-63 overexpression induces the formation of elongated lumenal ERmoxGFP blobs along ER tubules that overlap with mCherry-CLIMP-63. In contrast, upon mCherry-RTN4a overexpression, both ERmoxGFP and mCherry-RTN4a show highly periodic distributions that present minimal overlap (Fig. 3A).

**Fig. 3.**
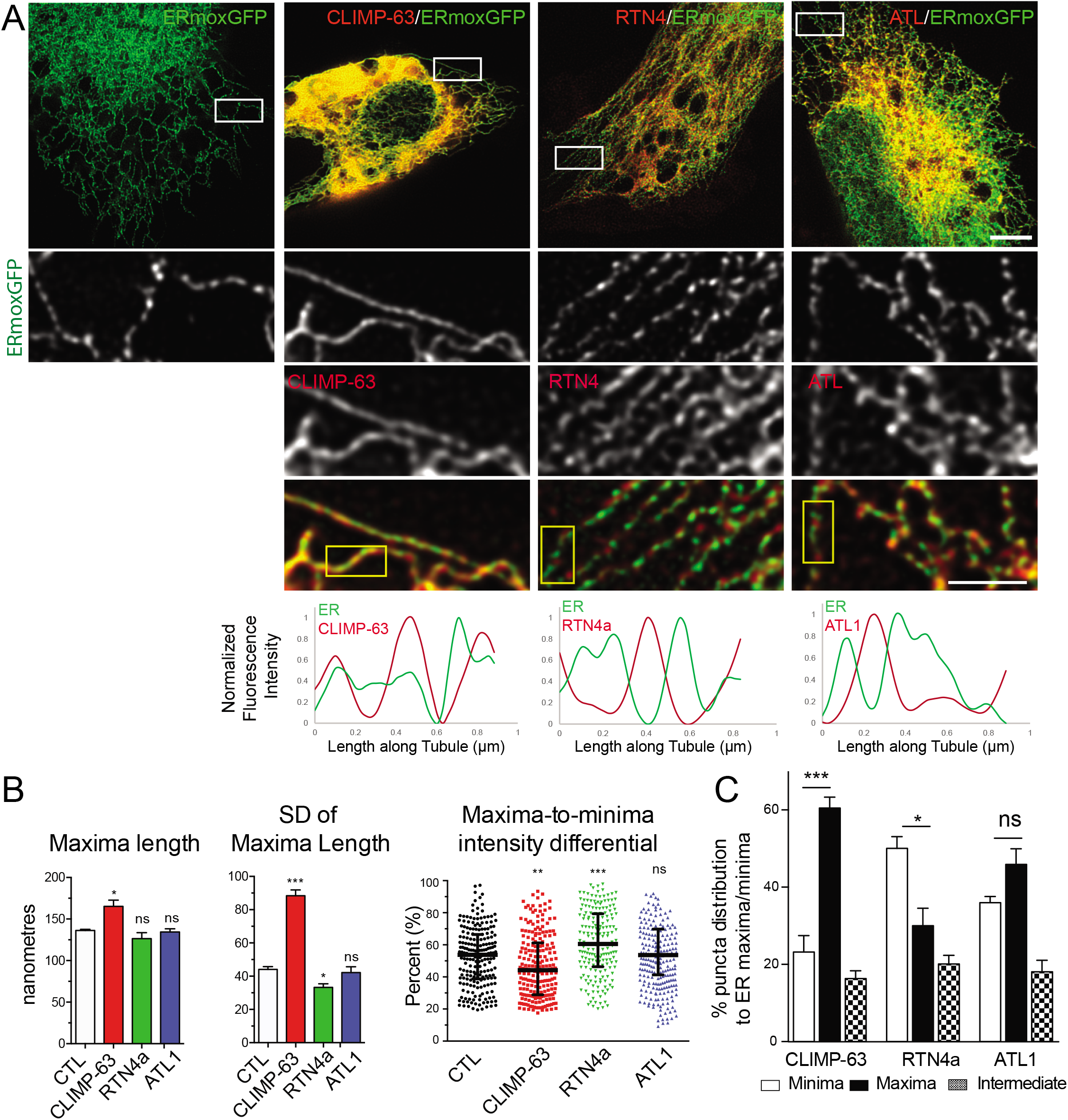
RTN4a and CLIMP-63 overexpression differentially impacts ER nanodomain periodicity. A) STED images of ERmoxGFP in HT-1080 cells transfected with ERmoxGFP or cotransfected with mCherry-CLIMP-63 (CLIMP-63), mCherry-reticulon4a (RTN4a) or mCherry-atlastin1 (ATL1). Peripheral ER regions (white boxes) are shown as zooms; line scans of selected tubules in these regions (yellow boxes) are shown with ERmoxGFP in green and ER shaping proteins in red. Scale bar: 5 μm (zooms: 2 μm). B) Peripheral ER tubule maxima length, variation of maxima length (SD), and maxima-to-minima intensity differential are shown for cells transfected with ERmoxGFP alone (CTL) or cotransfected with mCherry-CLIMP-63, mCherry-reticulon4a (RTN4a) or mCherry-atlastin1 (ATL1). Significance assessed by one-way ANOVA from three independent experiments (40 line scans/each repeat). Bar graphs show mean±SEM and scatter dot plots median with interquartile range. *, P < 0.05; **, P < 0.01; ***, P < 0.001; ns, not significant. C) Based on line scan analysis of peripheral ER tubules of HT-1080 cells cotransfected with mCherry-CLIMP-63, mCherry-reticulon (RTN4a) or mCherry-atlastin (ATL1), % localization of CLIMP-63, RTN4a and ATL1 puncta to minima or maxima of lumenal ERmoxGFP labeled tubules was quantified. Significance assessed by one-way ANOVA from three independent experiments (40 line scans/each repeat). Bar graphs show mean±SEM. *, P < 0.05; ***, P < 0.001; ns, not significant.

Line scan quantification of ERmoxGFP in peripheral ER tubules of HT-1080 cells shows that mCherry-CLIMP-63 transfection decreases lumenal periodicity. ERmoxGFP tubules present significantly increased maxima length, increased variation in maxima length and decreased maxima-to-minima intensity differentials. In contrast, overexpressed mCherry-RTN4a enhances the periodicity of ERmoxGFP, reducing variation of maxima length and increasing maxima-to-minima intensity differentials. mCherry-ATL1 overexpression does not impact maxima length, variation of maxima length or maxima-to-minima differentials (Fig. 3B). To quantify the extent of overlap of ERmoxGFP nanodomains with the different ER shaping proteins, we counted the number of ER shaping protein puncta localized to ERmoxGFP maxima or minima in line scans of individual peripheral ER tubules. mCherry-CLIMP-63 is significantly associated with ERmoxGFP maxima, mCherry-RTN4a with minima and mCherry-ATL1 shows no significant preference for maxima or minima (Fig. 3C). This suggests that RTN4a is segregated away from while CLIMP-63 is associated with lumenal ERmoxGFP filled nanodomains.

### RTN4a and CLIMP-63 regulate nanodomain heterogeneity in peripheral ER tubules

We then undertook to determine whether other ER markers present a similar peripheral ER tubule periodicity. As observed for ERmoxGFP, the peripheral reticular network of Sec61βGFP labeled tubules in live HT-1080 and COS-7 cells showed a highly periodic distribution by STED (Fig. 4A; Supp. Video 1). Similarly, GFP-calreticulin expressed at endogenous levels by CRISPR/Cas9 knock-in technology in U2OS cells (22) presents a tubular network by confocal and a highly periodic distribution by STED (Fig. 4A). 2D STED imaging of fixed cells expressing ERmoxGFP and Sec61βmRFP show distinct patterns of nanodomain enrichment for these two ER reporters along ER tubules (Fig. 4B).

**Fig. 4.**
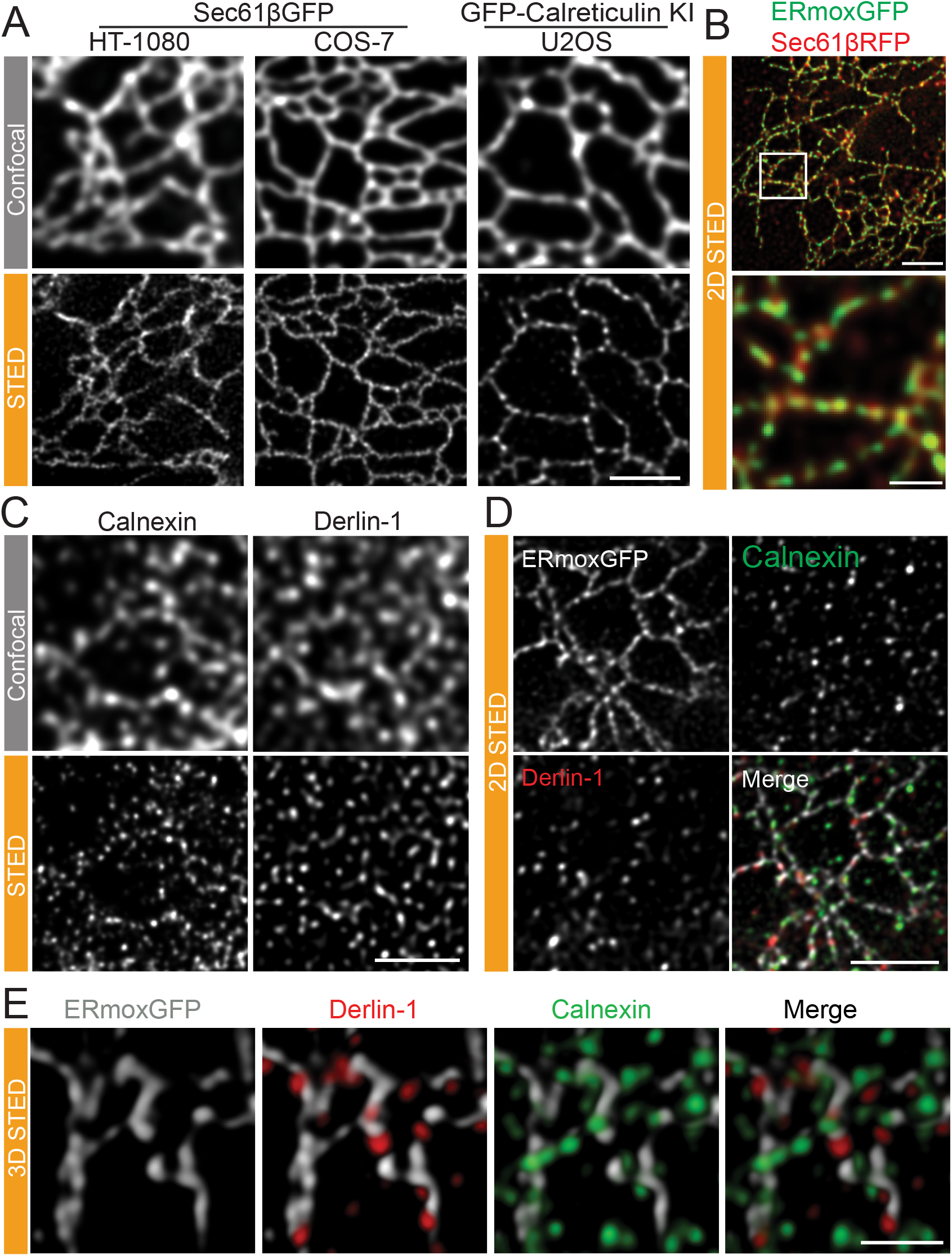
Periodic distribution of ER proteins along peripheral ER tubules. A) Representative confocal and STED images of live HT-1080 and COS-7 cells overexpressing Sec61βGFP or live knock-in U2OS cells expressing GFP-calreticulin at endogenous levels. Scale bar: 2 μm B) STED images of fixed HT-1080 cells expressing lumenal ERmoxGFP (green) and membrane Sec61βmRFP (red) show distinct periodicity of these two ER reporters along peripheral ER tubules. Scale bar: 2 μm; zoom: 0.5 μm. C) Untransfected HT-1080 cells labeled for derlin-1 or calnexin imaged by confocal and STED. Scale bar: 2 μm. D) Association of ER resident proteins derlin-1 and calnexin with ERmoxGFP labeled peripheral ER tubules in HT-1080 cells by 2D STED. Scale bar: 2 μm E) Association of ER resident proteins derlin-1 and calnexin with ERmoxGFP labeled peripheral ER tubules in HT-1080 cells by 3D STED. Scale bar: 1 μm.

We then extended our analysis to examine the distribution of the endogenous ER resident proteins, calnexin and derlin-1, involved in protein quality control and ERAD, respectively (25, 26). Antibody labeling of both calnexin and derlin-1 shows a reticular ER distribution by confocal microscopy in HT-1080 cells; by contrast, STED imaging of these endogenous ER proteins shows a highly punctate distribution (Fig. 4C). A similar punctate distribution for calnexin is observed in both ERmoxGFP transfected and untransfected cells (Supp. Fig. 1), and is therefore not a result of overexpression of the lumenal ERmoxGFP reporter. 2D STED images of fixed HT-1080 cells show that calnexin and derlin-1 puncta show minimal overlap and that many of these ER protein puncta align along ERmoxGFP labeled tubules (Fig. 4D). 3D STED analysis shows that the majority of calnexin and derlin-1 puncta are intercalated into the ERmoxGFP tubular network and that these two ER resident proteins show minimal overlap (Fig. 4E; Supp. Video 2), highlighting the nanodomain heterogeneity of peripheral ER tubules.

Quantitative line scan analysis of 2D STED imaged peripheral ER tubules shows the increased association of calnexin and derlin-1 puncta with ERmoxGFP minima and an equal distribution between minima and maxima of Sec61βGFP-labeled tubules (Fig. 5A,B). Similarly, manual counting of the distribution of calnexin or derlin-1 puncta in peripheral 3D STED ROIs (Fig. 4E) shows the clear localization of protein puncta to minima between ERmoxGFP labeled blobs (maxima) (Supp. Fig. 2A). Line scan analysis of the distribution of three other ER proteins, BiP/GRP78, Gp78, and Syntaxin-17 (Stx17), show a similar enriched distribution to ERmoxGFP minima along peripheral ER tubules (Supp. Fig. 2B). This suggests that ER tubule nanodomains that present increased accumulation of the lumenal ER reporter ERmoxGFP are segregated away from nanodomains enriched for ER resident protein complexes.

**Fig. 5.**
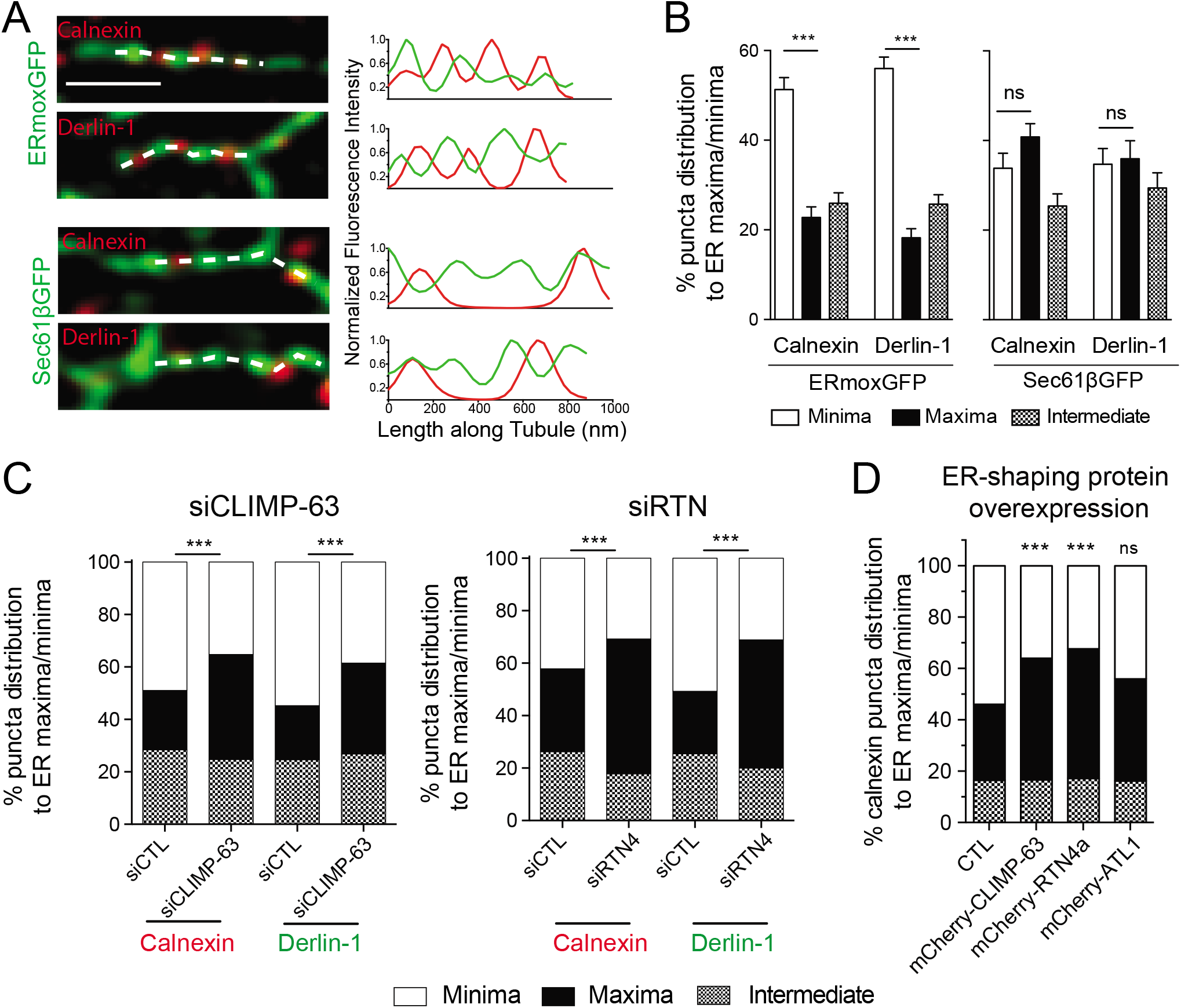
ER proteins are enriched in nanodomains depleted of lumenal ERmoxGFP. A) Representative merged images of single peripheral ER tubules expressing ERmoxGFP or Sec61βGFP labeled for calnexin or derlin-1. The dashed line indicates site of line scan analysis along tubule. Fluorescence intensities of ER reporter (green) and protein (red) from line scans are presented as graphs. Scale bar: 0.5 μm. B) Based on line scan analysis of peripheral ER tubules, % localization of calnexin and derlin-1 puncta to ERmoxGFP or Sec61βGFP maxima and minima was quantified. Values plotted are mean ± SEM from three independent experiments (40 tubules per repeat) with one-way ANOVA for significance. ***, P < 0.001; ns, not significant. C) Based on line scan analysis of peripheral ER tubules, % localization of calnexin and derlin-1 puncta to ERmoxGFP maxima and minima was quantified in cells transfected with siCTL, siCLIMP-63 or siRTN4. Significance assessed by χ2 test from three independent experiments (40 tubules per repeat). ***, P < 0.001. D) Based on line scan analysis of peripheral ER tubules, % localization of calnexin and derlin-1 puncta to ERmoxGFP maxima and minima was quantified in HT-1080 cells co-transfected with mCherry-CLIMP-63, mCherry-reticulon or mCherry-atlastin compared to control (CTL). Significance assessed by χ2 test from three independent experiments (40 tubules per repeat). ***, P < 0.001; ns, not significant.

We then tested if RTN4a and CLIMP-63 impact the distribution of ER resident proteins to ER tubule nanodomains. Upon RTN4 or CLIMP-63 knockdown or overexpression, calnexin or derlin-1 puncta no longer showed a preferential distribution to minima of lumenal ERmoxGFP but rather a balanced distribution to maxima and minima (Fig. 5C,D). ATL1 overexpression did not significantly impact the distribution of ER protein puncta to lumenal minima (Fig. 5D). This suggests that RTN4a and CLIMP-63 regulate lumenal domain length and organization and thereby impact overlap of lumenal and protein-enriched nanodomains along ER tubules.

### ER shaping proteins regulate lumenal nanodomain dynamics

To determine whether CLIMP-63 and reticulon impact the dynamics of ERmoxGFP lumenal nanodomains, we acquired high speed (40ms/frame) 2D STED time lapse image series of small ROIs encompassing individual peripheral ER tubules (Fig. 6a). Kymograms of 100 frames over 4 seconds show increased ERmoxGFP intensity in select locations along the tubule (Fig. 6b). This accumulation is indicative of lumenal filling of localized nanodomains that is reported as peaks when fluorescence intensity along the tubule is averaged over time (Fig. 6c). The coefficient of variation along tubule length of the normalized average fluorescent intensity over time provides a means to quantify the extent to which the lumenal ERmoxGFP reporter accumulates stably at defined locations along peripheral ER tubules (Fig. 6d). mCherry-CLIMP-63 overexpression disrupts the localized distribution of lumenal ER nanodomains while mCherry-RTN4a overexpression enhances the stability of these domains (Fig. 6A; Supp. Video 3). Conversely, siRTN4 knockdown showed a similar effect to CLIMP-63 overexpression, such that local stable accumulation of the lumenal reporter at discrete sites along ER tubules is no longer observed (Fig. 6B; Supp. Fig. 3; Supp. Video 4). CLIMP-63 knockdown did not affect lumenal ER nanodomain stability (Fig. 6B; Supp. Fig. 3; Supp. Video 4). The latter is consistent with the absence of an effect of CLIMP-63 siRNA on ERmoxGFP maxima length (Fig. 2C). Dynamic lumenal ERmoxGFP distribution along peripheral ER tubules therefore occurs within the framework of stable nanodomains whose ability to preferentially accumulate the lumenal ERmoxGFP reporter is regulated by reticulon and CLIMP-63.

**Fig. 6.**
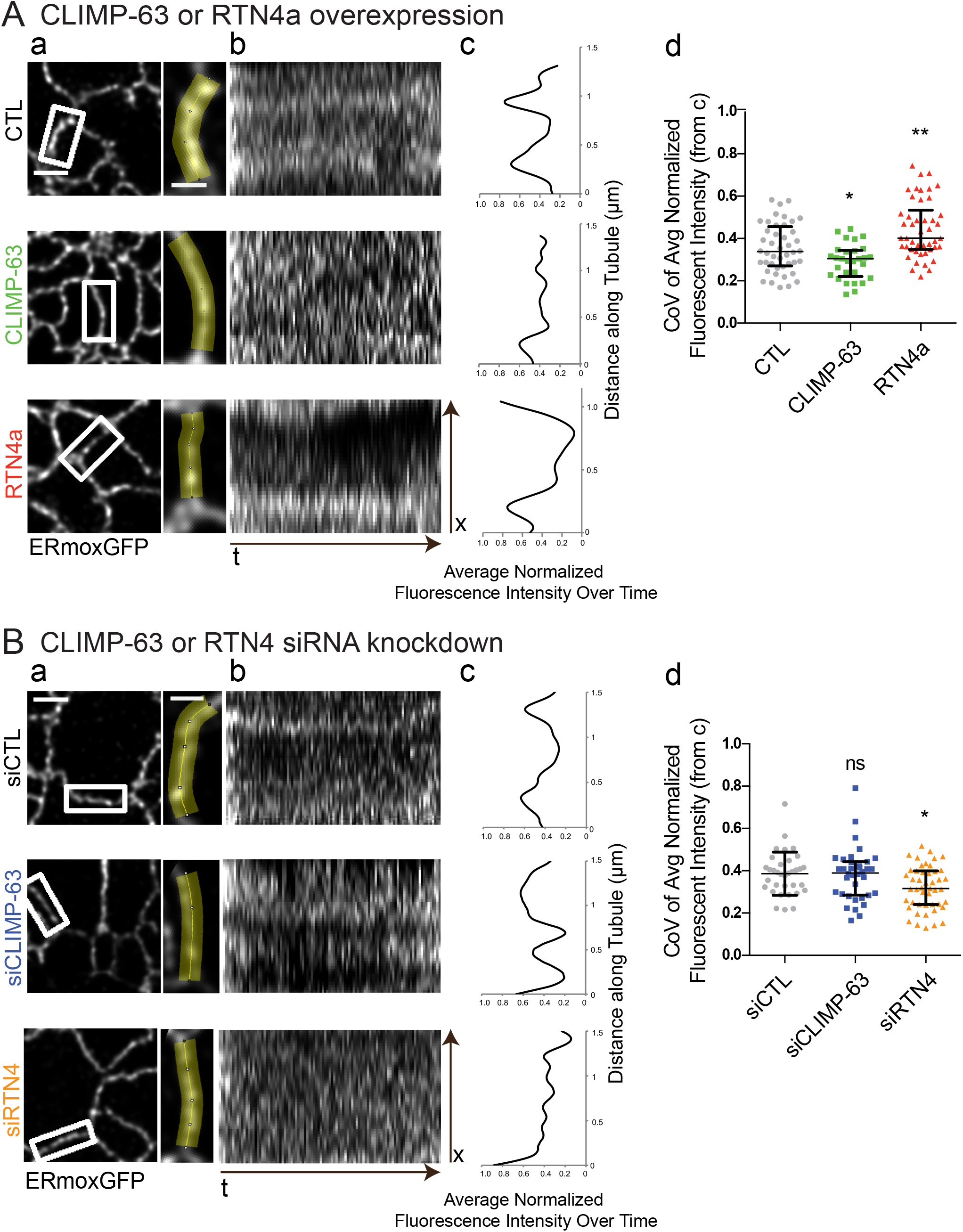
RTN4a and CLIMP-63 regulate dynamics of lumenal nanodomain compartmentalization along peripheral ER tubules. STED live cell imaging (40ms/frame over 4 seconds) of isolated ROIs of peripheral ERmoxGFP labeled tubules (a) was performed for COS-7 cells cotransfected with mCherry-CLIMP-63 or mCherry-RTN4a (A) or transfected with siCLIMP-63, siRTN4 or siCTL (B). Kymograms show the distribution of ERmoxGFP at specific sites along ER tubules over time (b). From plots of normalized average intensity over time (c), we determined the coefficient of variation (CoV) along tubule length as a measure of localized distribution of ERmoxGFP to distinct domains along peripheral ER tubules (d). Scatter dot plots show median with interquartile range from three independent experiments (20-50 tubules per condition) with one-way ANOVA for significance. *, P < 0.05; **, P < 0.01; ns, not significant.

## Discussion

Peripheral ER tubules are heterogenous, periodic structures composed of discrete nanodomains. STED imaging of lumenal (ERmoxGFP) or membrane (Sec61βGFP) ER reporters, of CRISPR/Cas9 knock-in calreticulin-GFP expressed at endogenous levels, and of various antibody labeled ER resident proteins, shows that ER proteins are localized to discrete puncta interspersed with lumenal domains along ER tubules. Periodic distribution of ER reporters along ER tubules can be observed in various publications studying ER using various super-resolution microscope approaches and EM (13, 27–32) and has more recently been reported by live cell GI-SIM (19). However, the nanodomain organization of ER tubules, and the mechanisms that underlie it have yet to be characterized. We show here that the ER shaping proteins RTN4a and CLIMP-63 regulate the size and stability of lumenal ER nanodomains and their overlap with resident ER proteins.

Single particle tracking has identified active lumenal flow within ER tubules (18). By high-speed 2D STED imaging, ER lumenal periodicities are highly dynamic and present rapid oscillations (Fig. 1; Supp. Video 1; Supp. Video 3, 4). We interpret ERmoxGFP periodicities to reflect regions or nanodomains along ER tubules that preferentially accumulate or are filled with this lumenal ER reporter. Localized distribution of these lumenal periodicities along ER tubules over time suggests that these sites (blobs/maxima) are propitious to accumulation of the lumenal reporter, relative to adjacent minima that limit lumenal reporter accumulation. RTN4 knockdown or CLIMP-63 overexpression reduces while RTN4a overexpression enhances the stability of sites of lumenal reporter accumulation along ER tubules. These data suggest that these two ER shaping proteins regulate lumenal domain spacing along peripheral ER tubules. Our data does not however report directly on diffusion of the lumenal ERmoxGFP reporter in ER tubules. The relationship between RTN4a and CLIMP-63 regulation of ER nanodomains and the slower diffusion of lumenal ER reporters in ER tubules relative to cytoplasm (33) or the contractions associated with nano-peristalsis along ER tubules (18) remains to be determined.

The reticulons cause local ER curvature (9) and CLIMP-63 maintains ER architecture by forming coiled-coil structures that maintain lumenal spacing (6, 34). Knockdown of CLIMP-63 has been reported to decrease the width of ER sheets from 45.5 nm to 27.9 nm (6). While CLIMP-63 is predominantly associated with ER sheets (6, 34), it is also expressed in peripheral ER tubules (13). Paralleling the macro level segregation of these two ER shaping proteins (6), RTN4a segregates away from and CLIMP-63 associates with lumenal ERmoxGFP nanodomains. Alternating RTN4a membrane constriction and CLIMP-63 spacer functions along ER tubules could explain the lumenal ER periodicity that we report here. Indeed, a recent study of lumenal mEmerald-KDEL dynamics in peripheral ER tubules attributed the periodic distribution of this reporter to constrictions and bulges along the tubule length (19), as observed by EM (13, 35–37). However, ER tubule width is below the 70-80 nm STED resolution obtained in this study and our data cannot therefore report on changes in ER tubule width. Further, the fact that ERmoxGFP minima overlap with ER protein densities (Fig. 5) argues that lumenal minima are not necessarily constrictions of the ER membrane itself. Mechanisms that control lumenal filling of peripheral ER nanodomains may be related not only to tubule width (i.e. constrictions vs bulges) but also to occlusion of lumenal space within the tubules by ER resident protein complexes.

Indeed, by STED, puncta of various ER proteins (calnexin, derlin-1, Gp78, BiP and Stx17) along ER tubules were closely associated with reticulon-associated lumenal ERmoxGFP minima and therefore segregated away from lumenal nanodomains. Overexpression or knockdown of either CLIMP-63 or RTN4a disrupted the enrichment of resident ER proteins in nanodomains depleted of the lumenal ERmoxGFP reporter. Simplistically, increased ER tubule lumenal filling due to CLIMP-63 overexpression or RTN4 knockdown could facilitate overlap between resident ER proteins and fluid components while, conversely, localized accumulation of overexpressed reticulon along ER tubules may sequester both protein and lumenal ER tubule components away from reticulon-enriched nanodomains. That CLIMP-63 knockdown had minimal effect on lumenal nanodomain stability and distribution argues that it is but one of multiple redundant mechanisms to define lumenal enriched ER nanodomains. Regulation of ER nanodomain organization by these ER shaping proteins in peripheral ER tubules parallels their roles in the formation of ER tubules and ER sheets. Reticulon and CLIMP-63 also contribute to nanohole formation in ER sheets (17) and, together, these studies suggest that ER shaping proteins play critical roles in the determination of ER morphology from the macro to the nano scale.

This study suggests that ER tubules are composed of protein-enriched nanodomains interspersed with fluid-filled nanodomains that may facilitate diffusion and exchange of small molecules, such as metabolites and enzyme substrates, as well as folding intermediates between protein complexes. The minimal overlap between the two endogenous ER markers studied, calnexin and derlin-1, suggests that multiple, discrete protein complexes exist within the ER. Further study is required to define how nanodomain organization and heterogeneity within the highly dynamic ER tubular network controls functional interaction between ER resident proteins, cargo proteins, enzymatic substrates and metabolites to enable ER quality control and cargo processing.

## Materials and Methods

### Antibodies, plasmids, and chemicals

ERmoxGFP was a gift from Dr. Erik Snapp (Albert Einstein College of Medicine, presently at Howard Hughes Medical Institute Janelia Research Campus, Virginia) (Addgene plasmid # 68072), mCherry-CLIMP-63 and Sec61βGFP from Dr. Gia Voeltz (University of Colorado, Boulder), Sec61βmRFP from Dr. Patrick Lajoie (University of Western Ontario, London, Ontario), and mCherry-RTN4a and mCherry-ATL1 from Dr. Tom Rapoport (Harvard University, Massachusetts) (Addgene plasmid #86683 and #86678, respectively). Anti-CLIMP-63 antibody (G1/296) was purchased from Enzo Life Sciences (Cat#: ALX-804-604-C100), rabbit anti-calnexin (Cat#: C4731), mouse anti-derlin-1 (Cat#: SAB4200148), rabbit anti-BiP (Cat#: G9043) and rabbit anti-Stx17 (Cat#: HPA001204) antibodies from Sigma, rabbit anti-RTN4/NOGO (Cat#: 10950-1-AP) and anti-Gp78 (Cat#: 16675-1-AP) antibodies from Proteintech, mouse anti-actin from Sigma (Cat#: A2228), goat anti-mouse HRP conjugate from Sigma (Cat#: AP308P), goat anti-rabbit HRP conjugate from Sigma (Cat#: AP307P) and goat serum from Thermo Fisher Scientific (Cat#: 16210-064). Goat anti-rabbit IgG (H+L) cross-adsorbed secondary antibody, Alexa Fluor 532 (Cat#: A-11009) and goat anti-mouse IgG (H+L) cross-adsorbed secondary antibody, Alexa Fluor 568 (Cat#: A-11004) from Thermo Fisher Scientific. siCLIMP-63 (Cat#: L-012755-01-0005), siRTN4 (L-010721-00-0005) and non-targeting Control siRNA (Cat#: D-001810-01-50) were from GE Life Sciences. 16% paraformaldehyde (Cat#: 15710) and 25% glutaraldehyde (Cat#: 16220) were from Electron Microscopy Sciences, USA. Other chemicals were from Sigma.

### Cell culture and transfection

HT-1080 cells were grown at 37°C with 5% CO_2_ in complete RPMI 1640 (Thermo Fisher Scientific, USA) containing 10% FBS (Thermo Fisher Scientific, USA) and 1% L-glutamine (Thermo Fisher Scientific, USA) unless otherwise stated. The U2OS knock-in cell line expressing GFP-calreticulin at endogenous levels generated with CRISPR/Cas technology was provided by Dr. Tom Rapoport (Harvard University, Massachusetts). COS-7 and U2OS cells were grown at 37°C with 5% CO_2_ in complete DMEM (Thermo Fisher Scientific, USA) containing 10% FBS and 1% L-glutamine. All cell lines were mycoplasma-free, tested routinely by PCR (Applied Biological Materials, Canada) and, as necessary, treated with BM Cyclin (Roche, Germany) to eliminate mycoplasma contamination.

Plasmids were transfected for 22 hours using Effectene (Qiagen, Germany) according to the manufacturer’s protocols. siRNA transfection was done with Lipofectamine 2000 Transfection Reagent (Thermo Fisher Scientific, USA) in Opti-MEM Reduced Serum Media (Thermo Fisher Scientific, USA) which was replaced with fresh complete RPMI 1640 after 5 hours; after a further 21 hours plasmids were transfected with Effectene and incubated for 22 hours before fixation. For live cell imaging, cells were grown in ibidi 8-well μ-slides with #1.5H (170 μm +/−5 μm) D 263 M Schott glass and transfected as described above. For HT-1080 cells, complete RPMI-1640 medium was replaced with RPMI-1640 medium (Sigma, USA) without sodium bicarbonate, supplemented with 1% L-glutamine, 10% FBS, 10% HEPES (Thermo Fisher Scientific, USA) before imaging. For COS-7 and U2OS cells, complete DMEM medium was replaced by FluoroBrite DMEM media (Thermo Fisher Scientific, USA) before imaging. Imaging was conducted at 37°C.

### Immunofluorescence labeling

Cells grown on #1.5H coverslips (Paul Marienfeld, Germany) were: 1) fixed with 3% paraformaldehyde with 0.2% glutaraldehyde in PBS at room temperature for 15 minutes and washed with PBS-CM (PBS supplemented with 0.1 mM CaCl2 and 1 mM MgCl_2_; two quick washes and then two 5 minute washes); 2) permeabilized with 0.2% Triton X-100 for 5 minutes then washed with PBS-CM as above; 3) quenched with 1mg/mL of NaBH_4_ (Sigma, USA) for 10 minutes and washed with PBS-CM; 4) blocked with 10% Goat Serum (Thermo Fisher Scientific, USA) and 1% bovine serum albumin (Sigma, USA) in PBS-CM for 1 hour; 5) incubated with primary antibodies in Antibody Buffer (1% BSA, 2% goat serum, 0.05% Triton-X100, 20X sodium/sodium citrate buffer in Milli-Q H_2_O) overnight at 4°C then washed with PBS-CM then three times for 5 minutes with Antibody Wash Buffer (20x SSC, 0.05% Triton-X100 in Milli-Q H_2_O); 6) incubated with secondary antibodies in Antibody Buffer for 1 h then washed with PBS-CM then six times for 10 minutes with Antibody Wash Buffer on a rocker; 7) rinsed with Milli-Q H_2_O and mounted with ProLong Diamond (Thermo Fisher Scientific, USA) and cured for 24-48 hours at room temperature.

### Confocal and STED microscopy

Confocal and STED imaging was performed with the 100X/1.4 Oil HC PL APO CS2 STED White objective of a Leica TCS SP8 3X STED microscope (Leica, Germany) equipped with a white light laser, HyD detectors and Leica Application Suite X (LAS X) software (*LSI IMAGING*, Life Sciences Institute, University of British Columbia). Time-gated fluorescence detection was used for STED to further improve lateral resolution. Live cell time lapse imaging of GFP-tagged reporters was performed using the 592 nm depletion laser on a 15,360 nm square ROI (0.8 second frame rate over 40 seconds) or a 3,840 nm square ROI (40 ms frame rate over 4 seconds) in the periphery of the cell. For double or triple labeled fixed samples, acquisition was done at a scan speed of 600Hz with a line average of 6. GFP was excited at 488 nm and depleted using the 592 nm depletion laser. Alexa Fluor 532 was excited at 528 nm, Alex Fluor 568 at 577 nm and mRFP at 584 nm and all three were depleted using the 660nm depletion laser. Sequential acquisition (in the order of AF568/AF532/GFP or mRFP/GFP) between frames (2D) or between stacks (3D) was used to avoid crosstalk. 3D STED images were acquired at a step size of 100nm. STED images were deconvolved using Huygens Professional software (Scientific Volume Imaging, The Netherlands) that was also used to determine the theoretical PSFs from 2D STED live and fixed images and from 3D STED fixed images. XY Full Width at Half Maximum (FWHM) values obtained from the theoretical PSFs for STED GFP images were 70 nm for 2D fixed and 78 nm for 2D live. For 3D fixed analysis FWHM was 126 nm for XY and 340 nm for Z.

### Quantification and statistical analysis

Line scan analysis of at least 40 peripheral ER tubules per sample was done using Leica LAS-X software. Spatially isolated peripheral tubules were selected for analysis based on the presence of a minimum of two protein puncta per tubule in the protein labeled channel. A histogram of normalized fluorescence intensity (scale of 0-1) along the line was exported for the ER reporter (ERmoxGFP or Sec61βGFP), and either calnexin, derlin-1, mCherry-CLIMP-63, mCherry-RTN4a or mCherry-ATL1 and displayed using GraphPad Prism (GraphPad Software, USA). Maximum and minimum fluorescence values were identified with a Java script. Blobs were defined as local maxima in the fluorescence signal and minima as troughs in fluorescence intensity, at least 20% below both adjacent maxima. FWHM of maxima was used to determine blob length. The percent decrease in fluorescence signal of all minima was calculated relative to the adjacent maximum of lower fluorescence. Protein (calnexin and derlin-1) puncta were localized to either maxima or minima and deemed intermediate if the puncta centre was greater than 20 nm from the peak of the maxima or minima. Quantification of protein puncta distribution to maxima and minima in 3D projections was scored manually in LAS-X. 3D STED volume rendering was done with LAS-X. Analysis of the live cell imaging (40ms/frame) was done with ImageJ/FIJI (38). General image processing (2D image exporting to tiff format and 3D volume rendering) and final image preparation (merging, zoom cropping and addition of the scale bar) for publication were performed using LAS-X and FIJI. Statistical analyses were done using Prism 6.0.

## Acknowledgements

Supported by the Canadian Institutes of Health Research (PJT-148698), the National Science and Engineering Research Council of Canada (RGPIN 227925-13) and the Canada Foundation for Innovation/British Columbia Knowledge Development Fund (LEF 30636) to IRN. GG is the recipient of a UBC Four Year Doctoral Fellowship and CZ the recipient of an NSERC Undergraduate Student Research Award. Imaging was performed at the UBC Life Sciences Institute Imaging Core Facility (*LSI IMAGING*). We thank Dr. Christopher Loewen (UBC) for helpful discussions.

## Author contributions

Author contributions: G. Gao: conceptualization, designing and performing experiments, formal analysis, validation, methodology, project administration, figure preparation, visualization, and writing (draft preparation, review, and editing). E. Liu and C. Zhu: image analysis, quantification and figure preparation. I. R. Nabi: conceptualization, funding acquisition, methodology, project administration, supervision, and writing (draft preparation, review, and editing).

## Supplemental Figure Legends

**Supp. Fig. 1.**
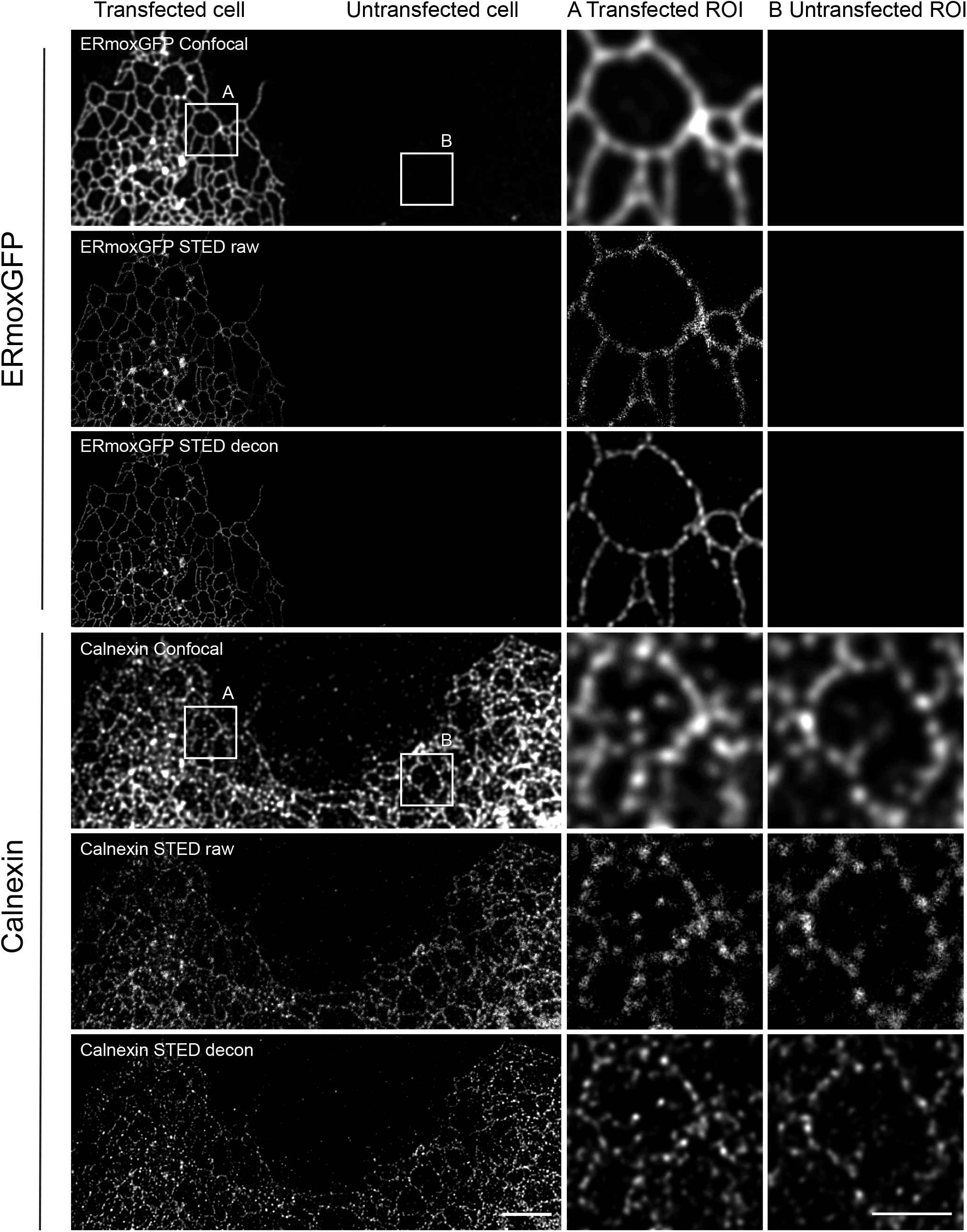
Calnexin exhibits a punctate distribution along peripheral ER tubules independent of ERmoxGFP transfection. Representative confocal and STED images of calnexin and ERmomxGFP in ERmoxGFP transfected and untransfected cells. The punctate distribution of calnexin is observed more readily by STED (raw and decon) compared to confocal in both (A) transfected and (B) untransfected cells. Scale bar: 5 μm; zooms: 2 μm.

**Supp. Fig. 2:**
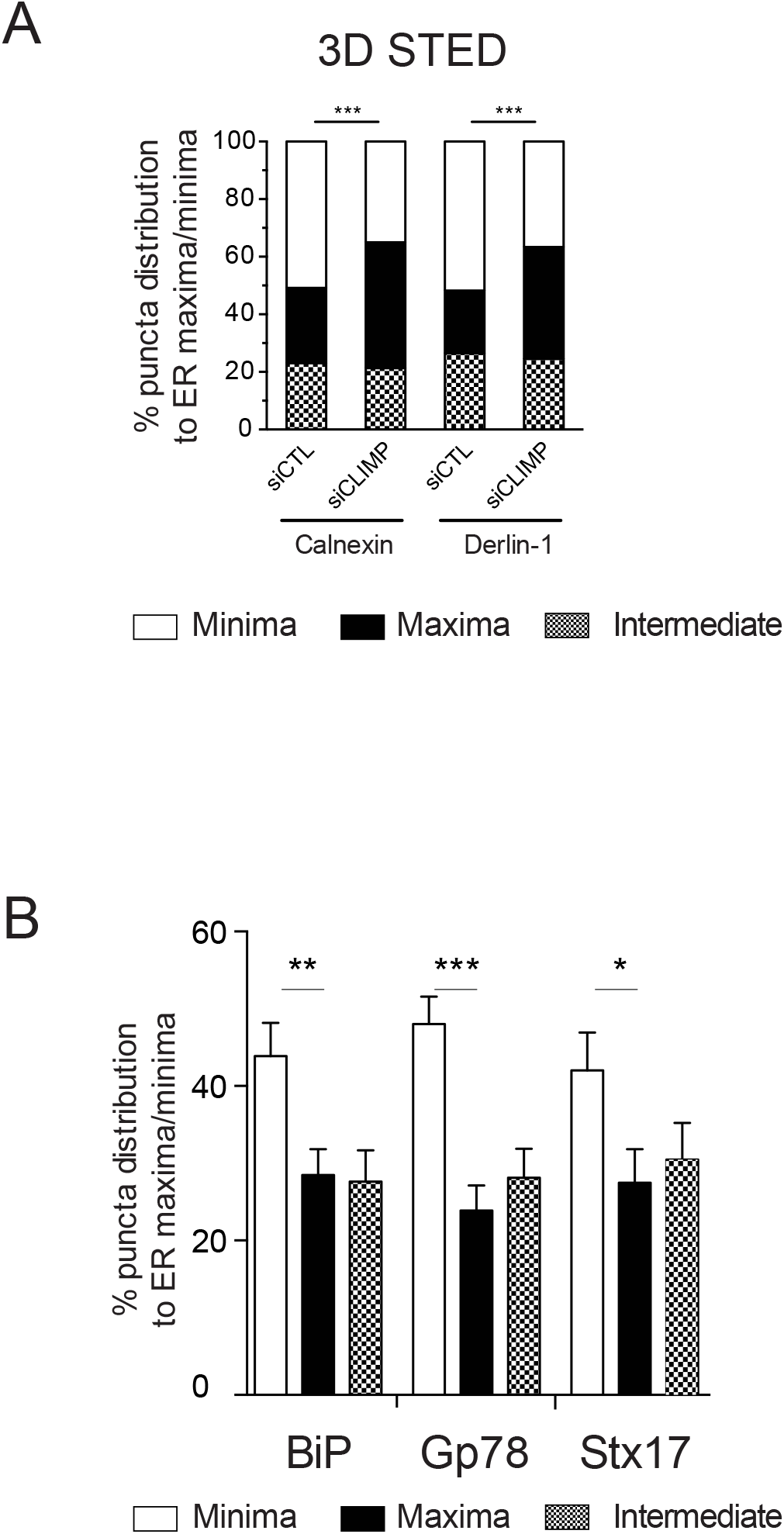
ER resident proteins are distributed to lumenal minima of peripheral ER tubules. (A) Quantification of three-dimensional localization of protein puncta to maxima and minima of peripheral ER tubules in siCTL or siCLIMP-63 HT-1080 cells by 3D STED, localization of calnexin and derlin-1 puncta to ERmoxGFP maxima and minima was quantified in siCTL or siCLIMP-63 HT-1080 cells. Significance assessed by χ2 test from at least 25 ROIs (2.5 um X 2.5um) from 10 3D stacks for each condition at two degrees of freedom in three independent experiments. ***, P < 0.001. (B) Based on line scan analysis of peripheral ER tubules imaged by 2D STED, localization of BiP. Gp78 and Stx17 puncta to ERmoxGFP maxima and minima was quantified. Significance assessed by one-way ANOVA from at least 120 line scans in three independent experiments. *, P < 0.05; **, P < 0.01; ***, P < 0.001.

**Supp. Fig. 3.**
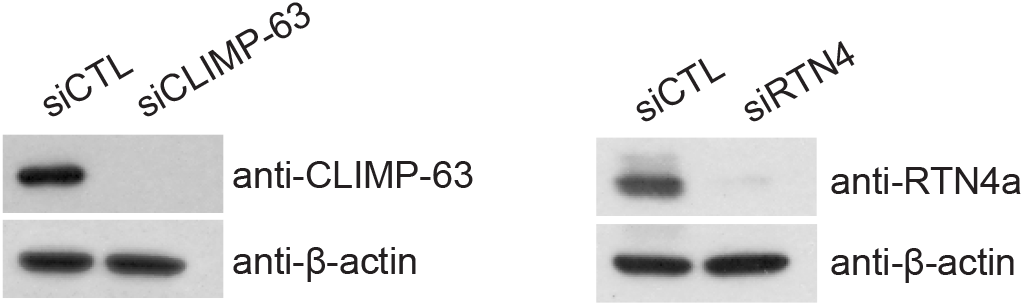
CLIMP-63 and reticulon knockdown by siRNA in COS-7 cells. Western blots of reticulon and CLIMP-63 siRNA knockdown in COS-7 cells. The blots were probed with anti-CLIMP-63, anti-RTN4a or anti-β-actin as a loading control.

## Supplemental Video Legends

Supp. Video 1: **Live cell STED imaging reveals ER periodicity.**

Live HT-1080 cells overexpressing ERmoxGFP or Sec61βGFP were imaged with confocal or STED at 37°C for 40 seconds at a temporal resolution of 0.8 s per frame. (a) confocal ERmoxGFP; (b) STED ERmoxGFP; (c) confocal Sec61βGFP; (d) STED Sec61βGFP. Video frame rate: ten frames/s.

Supp. Video 2: **3-channel 3D STED imaging of calnexin and derlin-1 puncta localized to ER tubule minima.**

ERmoxGFP expressing HT-1080 cells were fixed, labeled with calnexin and derlin-1 and imaged sequentially (derlin-1, calnexin and ERmoxGFP) with 3D STED (vortex donut enabled). Video frame rate: ten frames/s.

Supp. Video 3: **High-speed STED imaging of lumenal ER nanodomains upon RTN4a and CLIMP-63 overexpression.**

Live COS-7 cells were transfected with ERmoxGFP (CTL) or cotransfected with ERmoxGFP and either mCherry-CLIMP-63 (CLIMP-63 OX) or mCherry-RTN4a (RTN OX). ERmoxGFP was imaged with STED at 37°C over 4 seconds at a temporal resolution of 40 ms per frame. Arrows show sites of stable distribution of ERmoxGFP at specific sites along ER tubules over time, as determined by kymogram analysis (Fig. 6). Video frame rate: ten frames/s.

Supp. Video 4: **High-speed STED imaging of lumenal ER nanodomains upon RTN4 and CLIMP-63 knockdown.**

Live COS-7 cells were transfected with siCTL, siCLIMP-63 or siRTN4, as indicated, and then with ERmoxGFP. ERmoxGFP was imaged with STED at 37°C over 4 seconds at a temporal resolution of 40 ms per frame. Arrows show sites of stable distribution of ERmoxGFP at specific sites along ER tubules over time, as determined by kymogram analysis (Fig. 6). Video frame rate: ten frames/s.

